# Efficiently controlling for case-control imbalance and sample relatedness in large-scale genetic association studies

**DOI:** 10.1101/212357

**Authors:** Wei Zhou, Jonas B. Nielsen, Lars G. Fritsche, Rounak Dey, Maiken E. Gabrielsen, Brooke N. Wolford, Jonathon LeFaive, Peter VandeHaar, Sarah A. Gagliano, Aliya Gifford, Lisa A. Bastarache, Wei-Qi Wei, Joshua C. Denny, Maoxuan Lin, Kristian Hveem, Hyun Min Kang, Goncalo R. Abecasis, Cristen J. Willer, Seunggeun Lee

## Abstract

In genome-wide association studies (GWAS) for thousands of phenotypes in large biobanks, most binary traits have substantially fewer cases than controls. Both of the widely used approaches, linear mixed model and the recently proposed logistic mixed model, perform poorly – producing large type I error rates – in the analysis of phenotypes with unbalanced case-control ratios. Here we propose a scalable and accurate generalized mixed model association test that uses the saddlepoint approximation (SPA) to calibrate the distribution of score test statistics. This method, SAIGE, provides accurate p-values even when case-control ratios are extremely unbalanced. It utilizes state-of-art optimization strategies to reduce computational time and memory cost of generalized mixed model. The computation cost linearly depends on sample size, and hence can be applicable to GWAS for thousands of phenotypes by large biobanks. Through the analysis of UK Biobank data of 408,961 white British European-ancestry samples for >1400 binary phenotypes, we show that SAIGE can efficiently analyze large sample data, controlling for unbalanced case-control ratios and sample relatedness.

## Introduction

Decreases in genotyping cost allow for large biobanks to genotype all participants, enabling genome-wide scale phenome-wide association studies (PheWAS) in hundreds of thousands of samples. In a typical genome-wide PheWAS, GWAS for tens of million variants are performed for thousands of phenotypes constructed from Electronic Health Records (EHR) and/or survey questionnaires from participants in large cohorts^1,2^. For binary traits based on disease/condition status in PheWAS, cases are typically defined as individuals with specific International Classification of Disease (ICD) codes within the EHR. Controls are usually all participants without the same or other related conditions^1,2^. Due to the low prevalence of many conditions/diseases, case-control ratios are often unbalanced (case:control=1:10) or extremely unbalanced (case:control<1:100). The scale of data and the unbalanced nature of binary traits pose substantial challenges for genome-wide PheWAS in biobanks.

Population structure and relatedness are major confounders in genetic association studies and also need to be controlled in PheWAS. Linear mixed models (LMM) are widely used to account for these issues in GWAS for both binary and quantitative traits^3–8^. However, since LMM is not designed to analyze binary traits, it can have inflated type I error rates, especially in the presence of unbalanced case-control ratios. Recently, Chen, H. *et al*. have proposed to use logistic mixed models and developed a score test called the generalized mixed model association test (GMMAT)^9^. GMMAT assumes that score test statistics asymptotically follow a Gaussian distribution to estimate asymptotic p-values. Although GMMAT test statistics are more robust than the LMM based approaches, it can also suffer type I error rate inflation when case-control ratios are unbalanced, because unbalanced case-control ratios invalidate asymptotic assumptions of logistic regression^10^. In addition, since GMMAT requires O(*MN*^2^) computation and O(*N*^2^) memory space, where *M* is the number of genetic variants to be tested and *N* is the number of individuals, it cannot handle data with hundreds of thousands of samples.

Here, we propose a novel method to allow for analysis of very large samples, for binary traits with unbalanced case-control ratios, which also infers and accounts for sample relatedness. Our method, Scalable and Accurate Implementation of GEneralized mixed model (SAIGE), uses the saddlepoint approximation (SPA)^11,12^ to calibrate unbalanced case-control ratios in score tests based on logistic mixed models. Since SPA uses all the cumulants, and hence all the moments, it is more accurate than using the Gaussian distribution, which uses only the first two moments. Similar to BOLT-LMM^8^, the large sample size method for linear mixed-models, our method utilizes state-of-art optimization strategies, such as the preconditioned conjugate gradient (PCG) approach^13^ for solving linear systems for large cohorts without requiring a pre-computed genetic relationship matrix (GRM). The overall computation cost of this proposed method is O(*MN*), which is substantially lower than the computation cost of GMMAT^9^ and many popular LMM methods, such as GEMMA^7^. In addition, we reduce the memory use by compactly storing raw genotypes instead of calculating and storing the GRM.

We have demonstrated that SAIGE controls for the inflated type I error rates for binary traits with unbalanced case-control ratios in related samples through simulation and the UK Biobank data of 408,961 white British samples^14,15^. By evaluating its computation performance, we demonstrate the feasibility of SAIGE for large-scale PheWAS.

## RESULTS

### Overview of Methods

The SAIGE method contains two main steps: 1. Fitting the null logistic mixed model to estimate variance component and other model parameters. 2. Testing for association between each genetic variant and phenotypes by applying SPA to the score test statistics. Step 1 iteratively estimates the model parameters using the computational efficient average information restricted maximum likelihood (Al-REML) algorithm^16^, which is also used in GMMAT^9^. Several optimization strategies have been applied in step 1 to make fitting the null logistic mixed model practical for large data sets, such as the UK Biobank ^14,15^. First, the spectral decomposition has been replaced by the PCG to solve linear systems without inversing the *N* × *N* GRM^13^ (as in BOLT-LMM^8^). The PCG method iteratively finds solutions of the linear system in a computation and memory-efficient way. Thus, instead of requiring a pre-computed GRM, which costs a significant amount of time to calculate when sample sizes are large, SAIGE uses the raw genotypes as input. The computation time is about O(*M*_1_*N*) times the number of iterations for the conjugate gradient to converge, where *M*_1_ is a number of variants to be used for constructing GRM. Second, to further reduce the memory usage during the model fitting, the raw genotypes are stored in a binary vector and elements of GRM are calculated when needed rather than being stored, so the memory usage is *M*_1_*N/4* bytes (as in BOLT-LMM^8^ and GenABEL^17^). For example, for the UK Biobank data with *M*_1_ = 93,511 and *N* = 408,961 (white British participants), the memory usage drops from 669 Gigabytes(Gb) for storing the GRM with float numbers to 9.56 Gb for the raw genotypes in a binary vector.

After fitting the null logistic mixed model, the estimate of the random effects for each individual is obtained. The ratio of the variances of the score statistics with and without incorporating the variance components for the random effects is calculated using a subset of randomly selected genetic variants, similar to BOLT-LMM^8^ and GRAMMAR-Gamma^18^. This ratio has been previously suggested to be constant for score tests based on LMMs^18^. We have shown that the ratio is also approximately constant for all genetic variants with MAC ≥ 20 in the scenario of the logistic mixed models through analytic derivation and simulations **(Supplementary Notes and Supplementary Figure 1)**.

In step 2, for each variant, the variance ratio is used to calibrate the score statistic variance that does not incorporate variance components for random effects. Since GRM is no longer needed for this step, the computation time to obtain the score statistic for each variant is O(*N*). SAIGE next approximates the score test statistics using the SPA to obtain more accurate p-values than the normal distribution. A faster version of the SPA test, similar to the fastSPA method in the SPAtest R package that we recently developed^12^, is used to further improve the computation time, which exploits the sparsity in low frequency or rare variants to reduce the computation cost.

### Computation and Memory Cost

The key features of SAIGE compared to other existing methods are presented in **Table 1**, showing that SAIGE is the only mixed-model association method that is able to account for the unbalanced case-control ratios while remaining computationally practical for large data sets. To further evaluate the computational performance of SAIGE, we randomly sampled subsets from the 408,458 white British UK Biobank participants who are defined as either coronary artery disease (CAD) cases (31,355) or controls (377,103) based on the PheWAS Code 411^2,14,15^ followed by benchmarking association tests using SAIGE and other existing methods on 200,000 genetic markers randomly selected out of the 71 million with imputation info ≥ 0.3. The non-genetic covariates sex, birth year, and principal components 1 to 4 were adjusted in all tests. The log10 of the memory usage and projected computation time for testing the full set of 71 million genetic variants are plotted against the sample size as shown in **Supplementary Figure 2** and **Supplementary Table 1**. Although SAIGE and BOLT-LMM have the same order of computational complexity (**Table 1**), SAIGE was slower than BOLT-LMM across all sample sizes (ex. 517 vs 360 CPU hours when *N*=408,458). This is due to the fact that fitting logistic mixed model requires more iterative steps than linear mixed model, and applying SPA requires additional computation. SAIGE requires slightly less memory than BOLT-LMM (10 to 11 Gb when *N*=408,458) and the low memory usage makes both methods feasible for the large data set. In contrast, GMMAT and GEMMA requires substantially more computation time and memory usage. For example, when *N*=400,000, projected memory usages of both GMMAT and GEMMA are more than 600 Gb. The actual computation time and memory usage of association tests for the full UK Biobank data for CAD are given in **Table 1**. SAIGE required 517 CPU hours and 10.3 Gb memory to analyze 71 million variants that have imputation info ≥ 0.3 for 408,458 samples, which indicates that the analysis will be done in ~26 hours with 20 CPU cores.

**Table 1.** Comparison of different methods for GWAS with mixed effect models

|  |  | Method Features |  |  |  |  | Algorithm Complexity |  |  |  | Benchmarks for UK Biobank Data Coronary Artery Disease (PheCode 411) |  |
| --- | --- | --- | --- | --- | --- | --- | --- | --- | --- | --- | --- | --- |
|  |  | Does not require pre-computed genetic relationship matrix | Feasible for large sample sizes | Developed for binary traits | Accounts for unbalanced case-control ratio | Tests quantitative traits | Time complexity |  | Memory usage (Gbyte) |  | Time CPU hrs | Memory |
|  |  |  |  |  |  |  | Step 1 | Step 2 | Step 1 | Step 2 |  |  |
| Logistic mixed model | SAIGE | ✓ | ✓ | ✓ | ✓ | ✓ | $O(PM_1N^{1.5})^*$ | $O(MN)$ | $M_1N/4$ | $N$ | 517 | 10.3G |
| | GMMAT | | | ✓ | | ✓ | $O(PN^3)$ | $O(MN^2)$ | $F N^2$ | $F N^2$ | NA | NA |
| Linear mixed model | BOLT-LMM | ✓ | ✓ | | | ✓ | $O(PM_1N^{1.5})^*$ | $O(MN)$ | $M_1N/4$ | $N$ | 360 | 10.9G |
| | GEMMA | | | | | ✓ | $O(N^3)$ | $O(MN^2)$ | $F N^2$ | $FN^2$ | NA | NA |

N: number of samples
P: number of iterations required to reach convergence
$M_1$ : number of markers used to construct the kinship matrix;
M: total number of markers to be tested
F: Byte for floating number
\* Number of iterations in PCG is assumed as $O(N^{0.5})^8$

### Association analysis of binary traits in UK Biobank data

We applied SAIGE to several randomly selected binary traits defined by the PheWAS Codes (PheCode) of UK Biobank^2,14,15^ and compared the association results with those obtained from the method based on linear mixed models, BOLT-LMM^8^, and SAIGE without the saddlepoint approximation (SAIGE-NoSPA), which is asymptotically equivalent to GMMAT^9^. Due to computation and memory cost, the current GMMAT method cannot analyze the UK Biobank data. We restrict our analysis to markers directly genotyped or imputed by the Haplotype Reference Consortium (HRC)^19^ panel due to quality control issues of non-HRC markers reported by the UK BioBank. Approximately 28 million markers with minor allele counts (MAC) ≥ 20 and imputation info score > 0.3 were used in the analysis. Among 408,961 white British participants in the UK Biobank, 132,179 have at least one up to the third degree relative among the genotyped individuals^14,15^. We used 93,511 high quality genotyped variants to construct the GRM. In the UK Biobank data, most binary phenotypes based on PheCodes (1,431 out of 1,688; 84.8%) have case-control ratio lower than 1:100 **(Supplementary Figure 3)** and would likely demonstrate problematic inflation of association test statistics without SPA.

Association results of four exemplary binary traits that have various case-control ratios are plotted in Manhattan plots shown in **Figure 1** and in the quantile-quantile **(QQ)** plots stratified by minor allele frequency (MAF) shown in **Figure 2**. The four binary traits are coronary artery disease (PheCode 411) with 31,355 cases and 377,103 controls (1:12), colorectal cancer (PheCode 153) with 4,562 cases and 382,756 controls (1:84), glaucoma (PheCode 365) with 4,462 cases and 397,761 controls (1:89), and thyroid cancer (PheCode 193) with 358 cases and 407,399 controls (1:1138). In the Manhattan plots in **Figure 1**, each locus that contains any variant with p-value < 5×10^−8^ is highlighted as blue or green to indicate whether this locus has been reported by previous studies or not. **Supplementary Table 2** presents the number of all significant loci and those that have not been previously reported by each method for each trait and **Supplementary Table 3** lists all significant loci identified by SAIGE.

**Figure 1.**
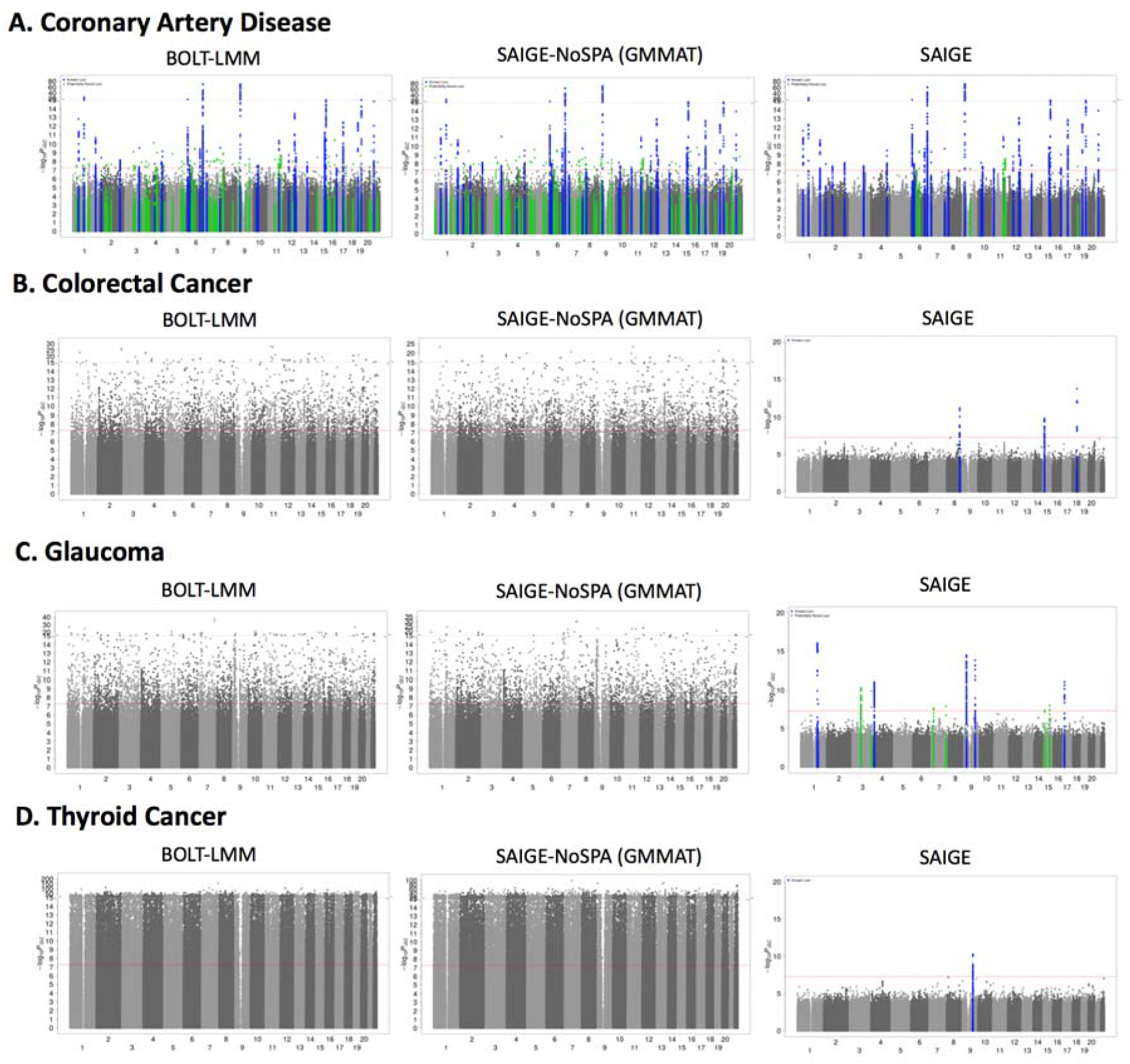
Manhattan plots of association p values resulting from SAIGE, SAIGE-NoSPA(asymptotically equivalent to GMMAT) and BOLT-LMM for A. coronary artery disease (PheCode 411, case:control = 1:12), B. colorectal cancer (PheCode 153, case:control = 1:84), C. glaucoma (PheCode 365, case: control = 1:89), and D. thyroid cancer (PheCode 193, case:control=1:1138). Blue: loci with association p-value < 5×10^−8^, which have been previously reported, Green: loci that have association p-value < 5×10^−8^ and have not been reported before. Since results from SAIGE-noSPA and BOLT-LMM contain many false positive signals for colorectal cancer, glaucoma, and thyroid cancer, the significant loci are not highlighted.

**Figure 2.**
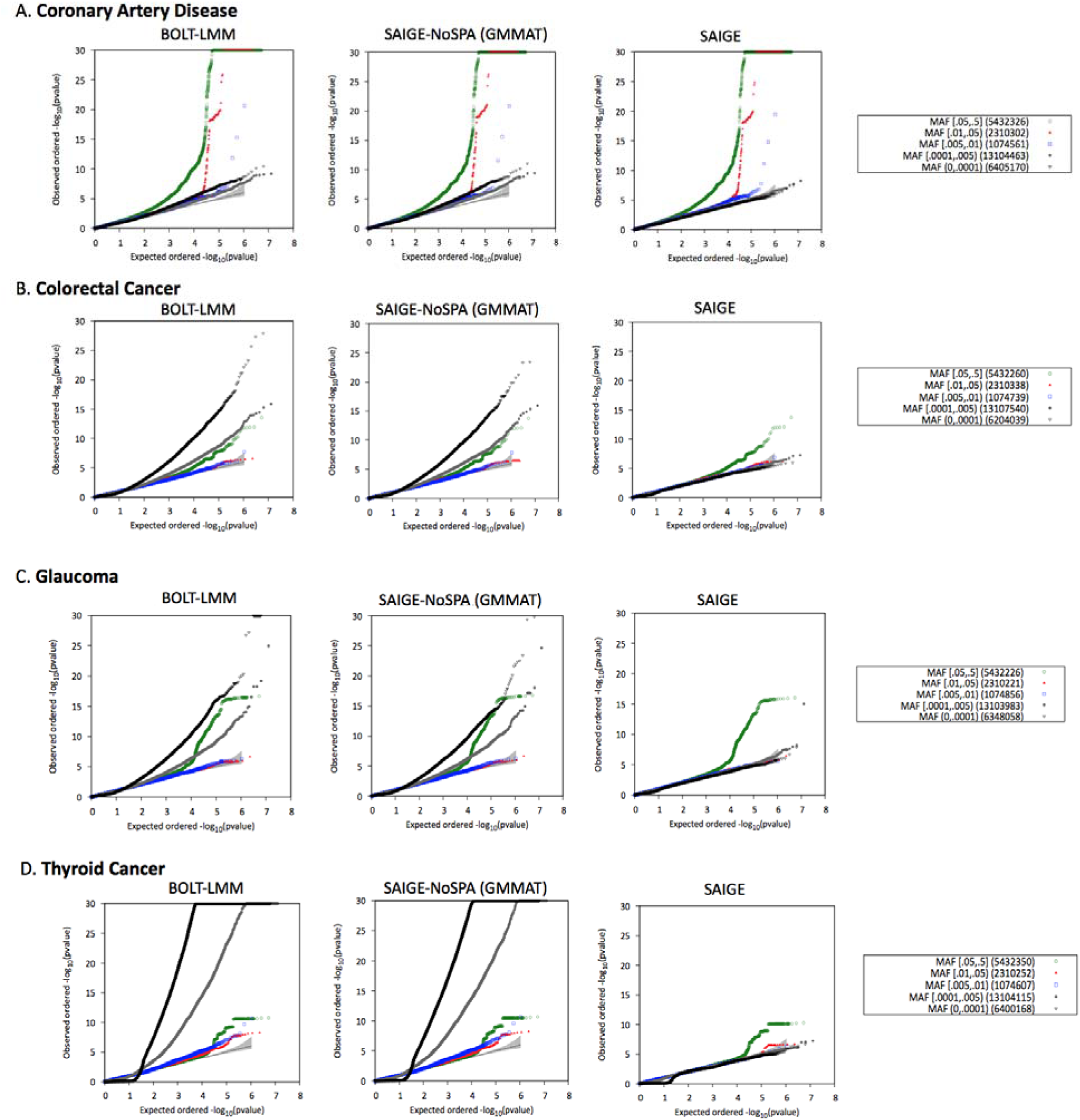
Quantile-quantile plots of association p-values resulting from SAIGE, SAIGE-NoSPA (asymptotically equivalent to GMMAT) and BOLT-LMM for A. coronary artery disease (PheCode 411, case: control = 1:12), B. colorectal cancer (PheCode 153, case: control = 1:84), C. glaucoma (PheCode 365, case: control = 1:89), and D. thyroid cancer (PheCode 193, case: control=1:1138).

Both Manhattan and QQ plots show BOLT-LMM and SAIGE-NoSPA have greatly inflated type I error rates. The inflation problem is more severe as case-control ratios become more unbalanced and the MAF of the tested variants decreases. The genomic inflation factors (λ) at the 0.001, 0.01 p-value percentiles are shown for several MAF categories in **Supplementary Table 4**. For the colorectal cancer GWAS which has case-control ratio 1:84, λ at the 0.001 p-value percentile is 1.68 and 1.71 for variants with MAF< 0.01 by SAIGE-NoSPA and BOLT-LMM, while λ is 0.99 by SAIGE. The inflation is even more severe for the test results by SAIGE-NoSPA and BOLT-LMM for the thyroid cancer, which has case-control ratio 1:1138, with the λ at the 0.001 p-value percentile around 4 to 5 for variants with MAF< 0.01 and all variants, respectively. With the unbalanced case-control ratio accounted for in SAIGE, the λ is again very close to 1.

We have generated summary statistics for all 1,403 PheCode-derived binary traits in 408,961 UK Biobank white British European-ancestry samples using SAIGE software and made them available in a public repository (see below for URL).

### Simulation Studies

We investigated the type I error control and power of two logistic mixed model approaches, SAIGE and GMMAT, and the linear mixed model method BOLT-LMM that computes mixed model association statistics under the infinitesimal and non-infinitesimal models through simulation studies. We followed the steps described on the Methods section to simulate genotypes for 1,000 families, each with 10 family members (N=10,000), based on the pedigree shown in **Supplementary Figure 4**.

### Type I error rates

The type I error rates for SAIGE, SAIGE-NoSPA, GMMAT, and BOLT-LMM have been evaluated based on the association tests performed on 10^9^ simulated genetic variants. The variants were simulated using the same MAF spectrum of the UK Biobank HRC imputation data with case-control ratio 1:99, 1:9, and 1:1. Two different values of variance component parameter and 2 were considered, which correspond to the liability scale heritability 0.23 and 0.38, respectively. The empirical type I error rates at the α = 5×10^−4^ and α = 5×10^−8^ are shown in the **Supplementary Table 5**. SAIGE-NoSPA, GMMAT, and BOLT-LMM have greatly inflated type I error rates when the case-control ratios are moderately or extremely unbalanced and slightly deflated type I error rates when the case-control ratios are balanced. This is expected as previous studies have suggested inflation of the score tests in the presence of the unbalanced case-control ratios and deflation in balanced studies^10,12^. We also observed that GMMAT score test statistics do not follow the normal distribution when MAF is low and case-control is unbalanced **(Supplementary Figure 5)**. Unlike GMMAT and BOLT-LMM, SAIGE has no inflation when case-control ratios are unbalanced. SAIGE also has no deflation when the case-control ratios are balanced.

To further investigate the type I error rates by MAF and case-control ratios, we carried out additional simulations. **Supplementary Figure 6** shows QQ plots of 1,000,000 rare variants (MAF = 0.005) with various case-control ratios (1:1, 1:9, and 1:99) and **Supplementary Figure 7** shows QQ plots of 1,000,000 variants with different MAF (0.005, 0.01, and 0.3) when case-control ratio was 1:99. Consistent to what has been observed in the real data study, GMMAT and SAIGE-NoSPA is more inflated for less frequent variants with more unbalanced case-control ratios. In contrast, SAIGE has successfully corrected this problem.

To evaluate whether SAIGE can control type I error rates in the presence of population stratification, we have simulated two subpopulations with Fst 0.013, which corresponds to the average Fst between Finnish and non-Finnish Europeans^20^. We assumed that subpopulations have different disease prevalences (0.01 for subpopulation 1 and 0.02 for subpopulation 2, 0.1 for subpopulation 1 and 0.2 for subpopulation 2, and 0.5 for subpopulation 1 and 0.4 for subpopulation 2). Both subpopulations have 1,000 families, each with 10 family members based on the pedigree shown in **Supplementary Figure 4**. Association tests were performed on 10 million simulated markers and the first four principle components were included as covariates **(Supplementary Figure 8)**. QQ plots **(Supplementary Figure 9)** show that the test statistics were well calibrated regardless of the variance component parameter *τ* and prevalence. This simulation result demonstrates that SAIGE produces well-calibrated p-values in the presence of population stratification.

### Power

Next, we evaluated empirical power. Since power simulation requires re-estimating a variance component parameter for each variant to test, to reduce computational burden, we used SAIGE-NoSPA instead of the original GMMAT software. Due to the inflated type I error rates of BOLT-LMM and GMMAT (and SAIGE-NoSPA), for a fair comparison, we estimated power at the test-specific empirical *α* levels that yield type I error rate α = 5×10^−8^ **(Supplementary Table 6). Supplementary Figure 10** shows the power curve by odds ratios for variants with MAF 0.05, 0.1 and 0.2 when *τ*=1. When the case-control ratio is balanced, the power of SAIGE, SAIGE-NoSPA and BOLT-LMM were nearly identical. For studies with moderately unbalanced case-control ratio (case:control=1:9), SAIGE has higher power than SAIGE-NoSPA and BOLT-LMM, which is due to very small empirical α for SAIGE-NoSPA and BOLT-LMM resulted from type I error inflation. The power gap is much larger when the case-control ratios are extremely unbalanced. Power results for *τ*=2 yielded the same conclusion regarding the methods comparison (data not shown).

Overall simulation studies show that SAIGE can control type I error rates even when case-control ratios are extremely unbalanced and can be more powerful than GMMAT and BOLT-LMM. In contrast, GMMAT and BOLT-LMM suffer type I error inflation, and the inflation is especially severe with low MAF and unbalanced case-control ratios.

### Code and data availability

SAIGE is implemented as an open-source R package available at https://github.com/weizhouUMICH/SAIGE/. The GWAS results for 1403 binary phenotypes with the PheCodes^2^ constructed based on ICD codes in UK Biobank using SAIGE are currently available for public download at https://www.dropbox.com/sh/wui4v8wsqiz78om/AAACfAJK54KtvnzSTAoaZTLma?dl=0.

**Supplementary Table 7** includes the phenotype information and URL links for downloading summary statistics, Q-Q plots, and Manhattan plots for the 1,403 phenotypes. We also display the results in the Michigan PheWeb http://pheweb.sph.umich.edu/UKBiobank, which consists of Manhattan plots, Q-Q plots, and regional association plots for each phenotype as well as the PheWAS plots for every genetic marker.

## DISCUSSION

In this paper, we have presented a method to perform the association tests for binary traits in large cohorts in the presence of sample relatedness, which provides accurate p-value estimates for even extremely unbalanced case-control settings (with a prevalence < 0.1%). The dramatic decrease of the genotyping cost over the last decade allows more and more large biobanks to genotype all of their participants followed by genome-wide PheWAS, in which GWASs are performed for all thousands of diseases/conditions characterized based on EHR and/or survey questionnaires to identify genetic risk factors across different phenotypes^1,2,21^. Several challenges exist for PheWAS studies by large cohorts. Statistically, inflated type I error rates caused by unbalanced case-control ratios and sample relatedness need to be corrected. Computationally, most of existing mixed model association methods are not feasible for large sample sizes. Our method, SAIGE, uses logistic mixed model to account for the sample relatedness and applies the saddle point approximation (SPA) to correct the inflation caused by the unbalanced case-control ratio in the score tests based on logistic mixed models.

SAIGE successfully corrects the inflation of type I error rates of low-frequency variants with binary traits that have unbalanced case-control ratios while also accounting for the relatedness among samples. Furthermore, our method uses several optimization strategies that are similar to those used by BOLT-LMM to improve its computational feasibility for large cohorts. For example, the preconditioned conjugate gradient algorithm is used to solve linear systems instead of the Cholesky decomposition method so that the time complexity for fitting the null logistic model is decreased from O(*N*^3^) to approximately O(*M*_1_*N*^1.5^), where *M*_1_ is the number of pruned markers used for estimating the genetic relationship matrix and the N is the sample size.

In the selection of genetic markers (*M*_1_) for estimating the kinship matrix and the variance component, trade off exists between computational cost and performance of adjusting for sample relatedness. Increasing the number of markers used for that step linearly increases the computation time and memory. On the other hand, using too few markers may not be sufficient to account for all subtle sample relatedness. For example, Yang *et al*. have shown that using a few thousand markers is not sufficient to yield correct type I error control^22^. In the UK Biobank data analysis, we used 93,511 independent, high quality genotyped variants, which were used by the UK Biobank data group to estimate the kinship coefficients between samples^15^. We carried out a sensitivity analysis by increasing the number of markers to 340,447 (Supplementary Section 2.3). Using more markers to estimate the kinship matrix for the UK Biobank data analysis produced generally similar association p-values but with lambdas closer to 1.

Using genome-wide genetic markers to adjust for sample relatedness tends to have the proximal contamination problem, which can reduce association test power^6,8,22,23^. To avoid it, the leave-one-chromosome-out (LOCO) scheme can be used. We implemented the LOCO option in SAIGE. A sensitivity analysis (Supplementary Section 1.2.5) on the four exemplary binary phenotypes in the UK Biobank suggested that the proximal contamination in GWAS for diseases with relatively low prevalence, such as thyroid cancer, glaucoma, and colorectal cancer, is not as substantial as for more common diseases, e.g. coronary artery disease.

Given the inflation of type I errors of linear mixed model for rare variants (MAF < 0.5%) with unbalanced case-control phenotypes, current GWAS studies address the problem by excluding rare variants from the analysis. However, this practice can lead to false negative results if associated rare variants are simply excluded rather than analyzed properly. For example, after using SAIGE to analyze rare variants in the UK Biobank, we identified a nonsense variant in *MYOC* (MAF = 0.14%) that was significantly associated with glaucoma. In our preliminary analysis of UK Biobank data of 1,283 non-sex specific phenotypes, we observed 1,609 genetic variants, including variants in the same locus, with minor allele frequency < 0.5% with SAIGE p-values < 5×10^−8^ (Supplementary Section 2.4). The method as implemented in SAIGE can control for type I error rates regardless of MAF and case-control ratios and will facilitate identification of rare disease-associated variants.

There are several limitations in SAIGE. First, the time for algorithm convergence may vary among phenotypes and study samples given different heritability levels and sample relatedness. Second, SAIGE has been observed to be slightly conservative when case-control ratios are extremely unbalanced **(Supplementary Table 5)**. Third, the accurate odds ratio estimation requires fitting the model under the alternative and is not computational efficient. Similar to several other mixed model methods^3,8,18^, SAIGE estimates odds ratios for genetic markers using the parameter estimates from the null model. Fourth, SAIGE assumes that the effect sizes of genetic markers are normally distributed with a mean of zero and standard deviation of one, which follows an infinitesimal architecture. With this assumption, SAIGE may sacrifice power to detect genetic signals whose genetic architecture is non-infinitesimal. Last, the variance component estimates 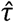 from SAIGE is biased and hence it should not be used to estimate the heritability (Supplementary Section 2.1). This is because SAIGE uses penalized quasi-likelihood (PQL) to estimate 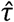. However, as shown in our simulation studies and elsewhere^9^, PQL-based approaches works well to adjust for sample relatedness. In future, we plan to extend the current single variant test to gene- or region-based multiple variant test to improve power for identifying disease susceptibility rare variants.

With the emergence of large-scale biobank, PheWAS will be an important tool to identify genetic components of complex traits. Here we describe a scalable and accurate method, SAIGE, for the analysis of binary phenotypes in genome-wide PheWAS. Currently, SAIGE is the only available approach to adjust for both case-control imbalance and family relatedness, which are commonly observed in PheWAS datasets. In addition, the optimization approaches used in SAIGE make it scalable for the current largest (UK Biobank) and future much larger datasets. Through simulation and real data analysis, we have demonstrated that our method can efficiently analyze a dataset with 400,000 samples and adjust for type I error rates even when the case-control ratios are extremely unbalanced. Our method will provide an accurate and scalable solution for large scale biobank data analysis and ultimately contribute to identify genetic mechanism of complex diseases.

## METHODS

### Generalized linear mixed model for binary traits

In a case-control study with sample size *N*, we denote the status of the *ith* individual using *y*_i_ = 1 or 0 for being a case or a control. Let the 1 × (1 + *p*) vector *X_i_* represent *p* covariates including the intercept and *G_i_* represent the allele counts (0,1 or 2) for the variant to test. The logistic mixed model can be written as

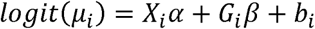

where *μ_i_* = *P*(*y*_i_ = 1 | *X_i_, G_i_, b_i_*) is the probability for the *ith* individual being a case given the covariates and genotypes as well as the random effect, which is denoted as *b_i_*. The random effect *b_i_* is assumed to be distributed as *N*(0, *τψ*), where *ψ* is an *N* × *N* genetic relationship matrix (GRM) and *τ* is the additive genetic variance. The *α* is a (1 + *p*) × 1 coefficient vector of fixed effects and *β* is a coefficient of the genetic effect.

### Estimate variance component and other model parameters (Step 1)

To fit the null model, *logit*(*μ*_*i*0_) = *X_i_α + b_i_*, penalized quasi-likelihood (PQL) method^24^ and the AI-REML algorithm^16^ are used to iteratively estimate 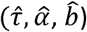. At iteration *k*, let 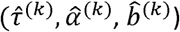 be estimated 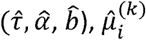 the estimated mean of *y_i_* 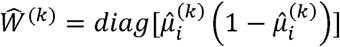 and 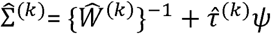 be an *n* × *n* matrix of the variance of working vector 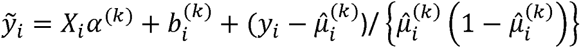. To obtain log quasi-likelihood and average information at each iteration, the current GMMAT approach calculates the inverse of 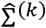. Since it is computationally too expensive for large *N*, we use the preconditioned conjugate gradient (PCG)^13,25^, which allows calculating quasi-likelihood and average information without calculating 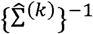. (See Supplementary for details). PCG is a numerical method to find solutions of linear system. It is particularly useful when the system is very large. BOLT-LMM^8^ successfully used it to estimate variance component in linear mixed model.

A score test statistics for *H_o_: β* = 0 is 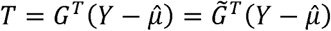 where *G* and *Y* are *N* × 1 genotype and phenotype vectors, respectively, and 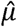 is the estimated mean of Y under the null hypothesis, and 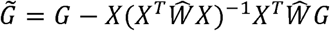 is the covariate adjusted genotype vector. The variance of *T*, Var(*T*) 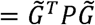, where 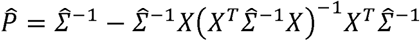. For each variant, given *P̂* calculation of Var(*T*) requires O(*N*^2^) computation. In addition, since our approach does not calculate 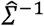, and hence *P̂*, obtaining Var(*T*) requires applying PCG for each variant, which can be computationally very expensive. To reduce computation cost, we use the same approximation approach used in BOLT-LMM and GRAMMAR-GAMMAR^18^, in which we estimate a variance of *T* with assuming that true random effect *b* is given, and then calculate ratio between these two variance. Suppose 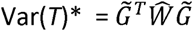, which is a variance estimate of T assuming *b̂* is given. Let *r* = Var(*T*)/Var(*T*)* ratio of these two different types of variance estimates. In Supplementary materials, we have shown that the ratio is approximately constant for all variants. Using this fact, we can estimate *r* using a relatively small number of variants. In all the numerical studies in this paper, we used 30 variants to estimate *r*.

### Score test with SPA (Step 2)

Suppose *r̂* is the estimated ratio (i.e. r) in Step 1. Now the variance adjusted test statistics is

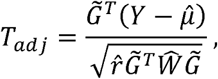

which has mean zero and variance 1 under the null hypothesis. The computation of *T_adj_* requires O(*N*) computation. The traditional score tests assume that *T* (and hence *T_adj_)* asymptotically follows a Gaussian distribution under *H_o_: β* = 0, which is using only the first two moments (mean and variance). When the case-control ratios are unbalanced and variants have low MAC, the underlying distribution of *T_adj_* can be substantially different from Gaussian distribution. To obtain accurate p-values, we use Saddlepoint approximation method (SPA)^11,12,26^, which approximates distribution using the entire cumulant generating function (CGF). A fast version of SPA (fastSPA)^12^ has recently been developed and applied to PheWAS, and provides accurate p-values even when case-control ratios are extremely unbalanced (ex. case:control=1:600).

To apply fastSPA to *T_adj_* we need to obtain CGF of T_adj_ first. To do this, we use the fact that given *b̂, T_adj_* is a weighted sum of independent Bernoulli random variables. The approximated cumulant generating function is

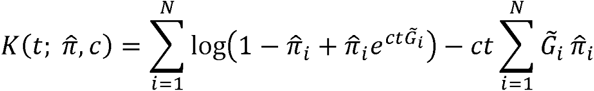

where the constant c=Var*(T)^−1/2^. Let *K′*(*t*) and *K″*(*t*) are first and second derivatives of K with respect to t. To calculate the probability that *T_adj_* < *q*, where q is an observed test statistic, we use the following formula^11^

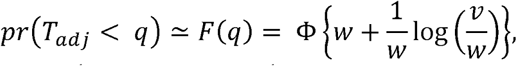

where 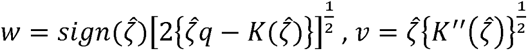 and 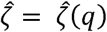 is the solution of the equation 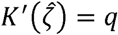. As fastSPA^12^, we exploit the sparsity of genotype vector when MAF of variants are low. In addition, since normal approximation works well when the test statistic is close to the mean, we use the normal distribution when the test statistic is within two standard deviation of the mean.

### Data simulation

We carried out a series of simulations to evaluate and compare the performance of SAIGE to GMMAT. We randomly simulated a set of 1,000,000 base-pair "pseudo" sequences, in which variants are independent to each other. Alleles for each variant were randomly drawn from Binomial(n = 2, p = MAF). Then we performed the gene-dropping simulation^27^ using these sequences as founder haplotypes that were propagated through the pedigree of 10 family members shown in **Supplementary Figure 4**. Binary phenotypes were generated from the following logistic mixed model

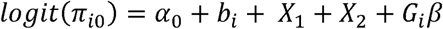

where *G_i_*, is the genotype value, *ß* is the genetic log odds ratio, *b_i_* is the random effect simulated from *N* (0,*τψ*) with *τ* = 1. Two covariates, X_1_ and X_2_, were simulated from Bernoulli(0.5) and N(0,1), respectively. The intercept *α*_0_ was determined by given prevalence (i.e. case-control ratios).

To evaluate the type I error rates at genome-wide α=5×10^−8^, 10 million markers along with 100 sets of phenotypes with different random seeds for case-control ratios 1:99, 1:9, and 1:1 were simulated with *β* = 0. Given *τ* = 1 and 2, the liability scale heritability is 0.23 and 0.38, respectively^28^ (Supplementary Section 2.1). Association tests were performed on the 10 million genetic markers for each of the 100 sets of phenotypes using SAIGE, GMMAT, and BOLT-LMM, therefore in total 10^9^ tests were performed. To have a realistic MAF spectrum, MAFs were randomly sampled from the MAF spectrum in UK Biobank data **(Supplementary Figure 11)**. Additional type I error simulations were carried out for five different MAFs (0.005, 0.01, 0.05, 0.1 and 0.3) to evaluate type I error rates by MAFs and in the presence of population stratification (Supplementary Section 2.2).

For the power simulation, phenotypes were generated under the alternative hypothesis *β* ≠ 0. For each of the MAF 0.05 and 0.2, we simulated 1,000 datasets, and power was evaluated at test-specific empirical α, which yields nominal α=5×10^−8^. The empirical α was estimated from the previous type I error simulations. As the same as type I error simulations, three different case-control ratios (1:99, 1:9, and 1:1) were considered.

Note that since we evaluated the empirical type I error rates and power based on genetic variants that were simulated independently, the LD Score regression^29^ calibration and the leave-one-chromosome-out (LOCO) scheme were not applied in BOLT-LMM or SAIGE.

### Phenotype definition in UK Biobank

We used a previously published scheme to defined disease-specific binary phenotypes by combining hospital ICD-9 codes into hierarchical PheCodes, each representing a more or less specific disease group^2^. ICD-10 codes were mapped to PheCodes using a combination of available maps through the Unified Medical Language System(https://www.nlm.nih.gov/research/umls/) and other sources, string matching, and manual review. Study participants were labeled a PheCode if they had one or more of the PheCode-specific ICD codes. Cases were all study participants with the PheCode of interest and controls were all study participants without the PheCode of interest or any related PheCodes. Gender checks were performed, so PheCodes specific for one gender could not mistakenly be assigned to the other gender.

## Acknowledgements

This research has been conducted using the UK Biobank Resource under application number 24460. SL and RD were supported by NIH R01 HG008773. CJW was supported by NIH R35 HL135824. WZ was supported by the University of Michigan Rackham Predoctoral Fellowship. JBN was supported by the Danish Heart Foundation and the Lundbeck Foundation. JCD, AG, LB, and WQW was supported by NIH R01 LM010685 and U2C OD023196.

## Author contributions

W.Z. C.W. and S.L. designed experiments. W.Z. and S.L. performed experiments. J.B., L.G.F., A.G., L.A.B, W-Q. W, J.C.D constructed phenotypes for the UK Biobank data. W.Z., J.L., S.A.G., H.M.K., C.W., S.L. and G.R.A. analyzed UK Biobank data. P.V. created the PheWeb. M.E.G. and K.H. provided data. W.Z., J.B., A.G., J.C.D., R.D., C.W. and S.L. wrote the manuscript.

## References

1. Bush, W. S., Oetjens, M. T. & Crawford, D. C. Unravelling the human genome-phenome relationship using phenome-wide association studies. Nat Rev Genet 17, 129–145 (2016).

2. Denny, J. C. et al. Systematic comparison of phenome-wide association study of electronic medical record data and genome-wide association study data. Nat Biotechnol 31, 1102–1110 (2013).

3. Kang, H. M. et al. Variance component model to account for sample structure in genome-wide association studies. Nat Genet 42, 348–354 (2010).

4. Zhang, Z. et al. Mixed linear model approach adapted for genome-wide association studies. Nat Genet 42, 355–360 (2010).

5. Yang, J., Lee, S. H., Goddard, M. E. & Visscher, P. M. GCTA: a tool for genome-wide complex trait analysis. Am J Hum Genet 88, 76–82 (2011).

6. Lippert, C. et al. FaST linear mixed models for genome-wide association studies. Nat Methods 8, 833–835 (2011).

7. Zhou, X. & Stephens, M. Genome-wide efficient mixed-model analysis for association studies. Nat Genet 44, 821–824 (2012).

8. Loh, P. R. et al. Efficient Bayesian mixed-model analysis increases association power in large cohorts. Nat Genet 47, 284–290 (2015).

9. Chen, H. et al. Control for Population Structure and Relatedness for Binary Traits in Genetic Association Studies via Logistic Mixed Models. Am J Hum Genet 98, 653–666 (2016).

10. Ma, C., Blackwell, T., Boehnke, M., Scott, L. J. & GoT2D investigators. Recommended Joint and Meta-Analysis Strategies for Case-Control Association Testing of Single Low-Count Variants. Genet. Epidemiol. 37, 539–550 (2013).

11. Kuonen, D. Saddlepoint approximations for distributions of quadratic forms in normal variables. Biometrika 4, 7 (1999).

12. Dey, R., Schmidt, E. M., Abecasis, G. R. & Lee, S. A fast and accurate algorithm to test for binary phenotypes and its application to PheWAS. Am. J. Hum. Genet. 101, 37–49 (2017).

13. E.F. Kaasschieter. Preconditioned conjugate gradients for solving singular systems. J. Comput. Appl. Math. 24, 265–275 (1988).

14. Sudlow, C. et al. UK Biobank: An Open Access Resource for Identifying the Causes of a Wide Range of Complex Diseases of Middle and Old Age. PLOS Med. 12, e1001779 (2015).

15. Bycroft, C. et al. Genome-wide genetic data on ~500,000 UK Biobank participants. bioRxiv 166298 (2017). doi:10.1101/166298

16. Gilmour, A. R., Thompson, R. & Cullis, B. R. Average Information REML: An Efficient Algorithm for Variance Parameter Estimation in Linear Mixed Models. Biometrics 51, 1440 (1995).

17. Aulchenko, Y. S., Ripke, S., Isaacs, A. & van Duijn, C. M. GenABEL: an R library for genome-wide association analysis. Bioinformatics 23, 1294–1296 (2007).

18. Svishcheva, G. R., Axenovich, T. I., Belonogova, N. M., van Duijn, C. M. & Aulchenko, Y. S. Rapid variance components–based method for whole-genome association analysis. Nat. Genet. 44, 1166–1170 (2012).

19. McCarthy, S. et al. A reference panel of 64,976 haplotypes for genotype imputation. Nat Genet 48, 1279–1283 (2016).

20. Nelis, M. et al. Genetic structure of Europeans: a view from the North-East. PLoS One 4, e5472 (2009).

21. Shameer, K. et al. A genome- and phenome-wide association study to identify genetic variants influencing platelet count and volume and their pleiotropic effects. Hum Genet 133, 95–109 (2014).

22. Yang, J., Zaitlen, N. A., Goddard, M. E., Visscher, P. M. & Price, A. L. Advantages and pitfalls in the application of mixed-model association methods. Nat. Genet. 46, 100–106 (2014).

23. Listgarten, J. et al. Improved linear mixed models for genome-wide association studies. Nat. Methods 9, 525–526 (2012).

24. Breslow, N. E. & Clayton, D. G. Approximate Inference in Generalized Linear Mixed Models. J. Am. Stat. Assoc. 88, 9 (1993).

25. Hestenes Eduard, M. R. and S. Methods of conjugate gradients for solving linear systems. 49, (NBS, 1952).

26. Imhof, J. P. Computing the Distribution of Quadratic Forms in Normal Variables. Biometrika Trust 48, 8 (1961).

27. Abecasis, G. R., Cherny, S. S., Cookson, W. O. & Cardon, L. R. Merlin–rapid analysis of dense genetic maps using sparse gene flow trees. Nat Genet 30, 97–101 (2002).

28. de Villemereuil, P., Schielzeth, H., Nakagawa, S. & Morrissey, M. General Methods for Evolutionary Quantitative Genetic Inference from Generalized Mixed Models. Genetics 204, 1281–1294 (2016).

29. Bulik-Sullivan, B. K. et al. LD Score regression distinguishes confounding from polygenicity in genome-wide association studies. Nat. Genet. 47, 291–295 (2015).

